# Costly induction of defense reduces plant growth and alters reproductive traits in mixed-mating *Datura stramonium*

**DOI:** 10.1101/2020.11.06.371989

**Authors:** Deidra J. Jacobsen

## Abstract

Herbivory shapes plant trait evolution by altering allocation to growth and defense in ways that affect plant reproduction and fitness. Initiation of these trade-offs may be particularly strong in juvenile plants with high phenotypic plasticity. Herbivory costs are often measured in terms of plant size or flower numbers, but other herbivore-induced floral changes can alter interactions with pollinators and have important implications for mating systems. In mixed-mating plants that can both self-fertilize and outcross, herbivory can maintain mating system variation if herbivore damage and defensive induction change a plant’s likelihood of selfing versus outcrossing. Here, I use mixed-mating *Datura stramonium* to evaluate how early defensive induction and herbivory result in trade-offs among plant defense, growth and reproduction. I used a 2×2 factorial manipulation of early chemical defense induction and season-long insecticide in the field. Growth costs of chemical induction were seen even before plants received damage, indicating an inherent cost of defense. Induction and herbivory changed multiple aspects of floral biology associated with a plant’s selfing or outcrossing rate. This including reduced floral allocation, earlier flowering, and reduced anther-stigma separation (herkogamy). Although these floral changes are associated with decreased attractiveness to pollinators, plants exposed to natural herbivory did not have decreased seed set. This is likely because their floral morphologies became more conducive to selfing (via reduced herkogamy). These vegetative and floral changes following damage and defensive induction can impact interactions among plants (by altering mating environment) and interactions with pollinators (via changes in floral allocation and floral phenology).

## Introduction

Plant-insect interactions can be used to understand how trophic interactions, abiotic/biotic variation, and phenotypic plasticity influence trait evolution. The costs of herbivory (including tissue loss and defensive induction) can result in trade-offs with plant growth or reproduction that can alter plant interactions with pollinators and decrease plant fitness (Karban 1993, Karban and Strauss 1993, Strauss *et al*. 2002, Stamp 2003, Barber *et al*. 2011). A trade-off between growth and defense on an macroevolutionary scale (i.e. examining traits among species) is well supported, with species that exhibit fast growth having low baseline levels of resistance (reviewed by Herms and Mattson 1992). However, within a species, this trade-off is less clear cut because of the complex way in which individual plants can respond to herbivory. Although defense can often come at the cost of decreased growth, these trade-offs can act to maximize reproductive success and fitness when under herbivore attack (Huot *et al*. 2014). Additionally, plants have a variety of ways to defend themselves against herbivory and different methods of defense may be more beneficial or less costly under different abiotic and biotic circumstances.

Plants employ both constitutive (always present) and induced (plastic) herbivore resistance but more work is needed to understand the costs of these defensive strategies and how these defenses affect plant fitness. Because induced defenses are produced or upregulated following a signal of herbivore attack, they are often considered a cost-saving strategy because they are only employed when needed (Karban and Baldwin 1997, Baldwin 1998). Costs of defense can come from either diversion of shared metabolic resources from growth or reproduction to defense or can be the result of less energetic resources left for defenses after production of plant defense (Mole 1994, Karban and Baldwin 1997, Machado *et al*. 2017, Züst and Agrawal 2017, Mesa and Dlugosch 2020). Regardless of mechanism, in order for an induced response to be a beneficial defensive strategy for the plant, it must increase plant fitness in the presence of herbivores (Karban and Baldwin 1997, Accamando and Cronin 2012). Whether the benefits of induced responses outweigh the costs ultimately depends on whether the allocation of resources to defense acts to preserve reproductive success and fitness.

For plants that rely on pollinators, changes in floral traits following herbivory and defensive induction are particularly consequential (Thompson and Johnson 2016). Reduced floral displays and decreased pollinator attraction can result in pollen limitation and lower seed set. When plants have a mixed-mating strategy, meaning they can both self-fertilize and outcross, the cost of reduced pollinator attraction following herbivory may be low if selfing is able to maintain seed set when pollinators are not adequately being recruited (Elle and Hare 2002), This balance between outcrossing (which minimizes inbreeding depression) and selfing (which provides reproductive assurance) can be influenced by the effects of herbivory on floral allocation (Ivey and Carr 2005, Steets *et al*. 2006, Campbell *et al*. 2013, Carr and Eubanks 2014). Because of this interaction between herbivory and selfing/outcrossing rate, herbivory has been proposed as a potential mechanism maintaining mixed-mating as a stable strategy despite theoretical models that predict the evolution of either nearly complete selfing or complete outcrossing for a species (Goodwillie *et al*. 2005, Ivey and Carr 2005, Bello-Bedoy and Nunez-Farfan 2010, Campbell and Kessler 2013).

Although floral display (flower size and number) is a clear contributor to pollinator attraction, herbivory may also alter floral rewards (such as pollen, nectar, or scent production) as well as flower shape and morphology (Kessler *et al*. 2011, Thompson and Johnson 2016). Herkogamy (the separation between the anthers and stigma) is a component of floral morphology that affects selfing. When the stigma is higher than the anthers, plants are unlikely to autonomously self-fertilize but when the anthers are at or above the level of the stigma, self-fertilization is likely. Therefore, herkogamy directly impacts whether a decrease in pollinator visitation is detrimental to seed set for mixed-mating species. Plastic changes in herkogamy could reduce the costs of induced responses by maintaining seed set via production of selfed seeds despite reduced attractiveness to pollinators and reductions in outcrossing (Kessler and Halitschke 2009).

Because of these complex interactions among herbivory, resource allocation, and mating system, understanding the costs associated with induced responses requires quantifying how herbivory and defensive induction alter plant growth, floral/reproductive allocation, and fitness over a plant’s lifespan. In the presence of natural pollinators and herbivores, the costs of induced responses can be overcome by benefits such as protection from future herbivory or increased survival. Therefore, examining these interactions in a field setting is necessary to understand how—and to what extent—induced responses translate into fitness costs or benefits.

In this study, I used a 2×2 factorial combination of early defensive induction and insecticide application in field-grown *Datura stramonium* L. (Solanaceae) to address how induction of plant defenses and herbivory alter plant growth and reproductive output/traits (including herkogamy) over the lifespan of this annual plant. This combination of treatments allowed me to test the costs of induction under natural and reduced levels of herbivore damage and collect lifetime fitness measurements. I found that herbivory and defensive induction resulted in long-lasting decreases in total growth and floral allocation. Following herbivory, floral morphology changed in a way that promoted selfing, which could maintain high seed set despite herbivore-driven decreases in floral allocation that negatively affect pollinator attraction.

## Materials and methods

### *Study system:* Datura stramonium

*Datura stramonium* is an annual plant commonly known as Jimson weed. *D. stramonium* is a mixed-mating species with self-fertilization rates between 81 and 100 percent (Motten and Antonovics 1992, Stone and Motten 2002) and is present throughout Central and North America (Bello-Bedoy and Nunez-Farfan 2011). Individual plants produce large flowers that are pollinated by nocturnal hawkmoths and generalist diurnal pollinators. Stigma position in *D. stramonium* generally ranges from 4 mm below to 12 mm above the anthers (Motten and Stone 2000). The stigma-anther separation is positively correlated with outcrossing rate. For flowers with anthers above the stigma (negative herkogamy), self-fertilization is highly likely, because pollen grains fall onto the receptive stigma (Motten and Antonovics 1992). Geitonogamy (self-pollination among different flowers on a plant) was shown to have little impact on outcrossing rates because of the small number of flowers open on a single plant simultaneously (Motten and Antonovics 1992). Additionally, because *D. stramonium* is rich in secondary compounds and has a variety of generalist and specialist herbivores, it is an ideal species with which to study plant defense (Castillo *et al*. 2014, Shonle and Bergelson 2000).

### Plant preparation prior to planting in field plot

Three populations of *Datura stramonium* were used for a common-garden field experiment in Bloomington, Indiana during 2014. The Indiana source populations used were: Wabash (“W”) (bank of Wabash River, lat 40.4160 N, long 86.9005 W), Highway 52 (“H”) (roadside, lat 40.4442 N, long 86.8695 W), and Applegate (“A”) (apple orchard, Trafalgar, IN lat 39.359 N, long 86.144 W). For each population, plants came from five maternal families to account for intrapopulation variation. Twelve plants from each family were used (N *=* 180 total). All seeds were surface sterilized for 30 minutes in a 50 percent bleach solution. Seeds were germinated 7 May 2014 on wet filter paper in a growth chamber (24 C, 14:10 light:dark cycle). Once plants had cotyledons, they were transferred to individual cell packs (200 cm^3^) containing MetroMix 360 until transplanting to the field on 7 June 2014.

### Early induction and insecticide treatments

The plastic induction of defense traits is often initiated by the production of jasmonic acid (JA), a phytohormone that coordinates many different defensive responses by allowing expression of defense genes (Eckardt 2008). Exogenous JA application can induce similar defensive chemicals and traits as those induced by herbivore damage in other Solanaceous species and was therefore used here to easily induce plant defenses in a controlled manner (Thaler *et al*. 1996). Plants from each maternal family were randomly assigned to one of four treatments in a 2×2 factorial design of jasmonic acid (JA) induction and insecticide application, ensuring that all maternal families and populations were equally represented in each treatment (N *=* 45 per treatment). Plants induced with an early JA treatment received two treatments (10 and 4 days before planting in the field) of 1mM exogenous JA to induce plant defenses (Thaler *et al*. 1996). JA dissolved in ethanol was diluted with dH2O to 1 mM and misted on plants to runoff. Control plants were sprayed with 1mM ethanol in dH2O and the two treatment groups were kept in separate rooms overnight to avoid volatile communication between plants in different treatment groups.

Insecticide-treated plants received a pre-treatment of Neem oil (Garden Safe, 4 tbsp/1 gallon H2O) two days prior to planting in the field. All surfaces of the plant were coated with insecticide. These insecticide-treated plants were re-sprayed weekly throughout the course of the experiment, whereas herbivore exposed plants were sprayed with only water. Treatments were applied near sunrise or sunset to avoid leaf burn.

### Field plot layout

Plants were distributed in the field to avoid insecticide drift onto control plants during spraying. This was done by splitting the field into 6 plots that each contained a group of insecticide-treated plants and a group of control (water spray) plants. Each alternating insecticide or control group contained 15 plants randomly assigned from the populations/families and JA treatments. Plants within a water or insecticide plot were 0.9 meters apart and each plot was 1.5-1.8 meters from another plot (Fig. S1). Plants were watered twice in the field at the time of transplanting to promote establishment.

### Growth, damage, and reproductive measurements

Plant size/growth was measured using weekly leaf counts from June to until flower production tapered off in early October 2014 (16 weeks). Weekly counts of damaged leaves were used to verify that the insecticide was effective at reducing the proportion of leaves receiving herbivore damage. Damage measurements were discontinued after 12 weeks, at which time all plants had flowered and set some matured fruits. While doing the damage estimates, any pollinators or herbivores seen on plants in the field were recorded. For herbivores that have clear feeding damage patterns or where the herbivore was observed feeding, the likely type of insect that caused the damage and the level of damage were recorded (e.x. flea beetles chew many tiny holes in the leaves).

Flowering date for each plant was recorded during daily surveys of flower number until complete freezing/death of plants following a hard freeze on 1 November 2014. To measure allocation to individual flowers, corolla length and herkogamy were measured the morning following evening corolla opening. Three early season flowers (first through third flowers) and three later season flowers (twenty-third through twenty-fifth flowers) were measured on each plant. These flowers were marked with wire and fruits were collected upon maturation for seed counts. The total number of flowers and mature fruit produced by each plant was recorded. Seeds were counted for the first and twenty-third flowers on each plant; if that fruit had been aborted, the fruit from the next flower opened was used for seed counts.

### Statistical analyses

I analyzed differences among the four treatments in herbivore damage and plant size over time using a repeated-measures mixed model implemented in PROC-MIXED in SAS 9.4 (SAS Institute, Cary NC). These repeated-measures mixed-effects models were used with either damage level (proportion of leaves damaged) or plant size (leaf numbers) as the response variable and fixed effects of: time/date, population, insecticide treatment (and its interaction with time), JA treatment (and its interaction with time), and the three-way interaction of time*insecticide treatment*JA treatment. Because each plant was re-measured weekly, date was included as a repeated factor with a first order autoregressive structure because individual plant sizes are expected to be correlated through time.

Because the repeated-measures analyses revealed variation over the course of the season in plant damage and size, additional mixed-effect models were used at each timepoint to determine the specific ways in which the JA induction and insecticide treatments affected plant size and damage. Population, insecticide treatment, JA treatment, and all interactions were used as fixed effects in separate models for plant size and plant damage. Random effects included maternal family, all interactions between fixed effects and family (to estimate variation between families), as well as block and interactions between block and the other random effects.

Mixed-effects models were also used to analyze reproductive traits for each plant. Separate analyses were used for each of the following response variables: flowering time, total flower number, flower size (early and late), herkogamy (early and late), total fruit number and seed set per fruit. Fixed effects were population, insecticide treatment, JA treatment, and all interactions. Random effects included maternal family, all interactions between fixed effects and family (to estimate variation among families), as well as block and interactions between block and the other random effects.

Box-cox power transformations were used for all vegetative floral and reproductive traits to correct for non-normality of residuals, except for flowering date (which was non-normal despite attempted corrections). Box-Cox-lambda (Box and Cox 1964) was calculated using PROC-TRANSREG. Because herkogamy values contained zero and negative numbers, all values were scaled by adding the lowest measurement for these traits, “5” (for early herkogamy) and “8” (for late herkogamy), to measured values prior to Box-Cox transformations and analyses. Similarly, for dates where the leaf damage estimates contained values of zero, each value was scaled by adding “1” to measured values prior to Box-Cox transformations and analysis (Table S1). Variance attributed to plant family was non-significant but was included as a random effect to account for experimental design.

To account for potential type I error caused by an unbalanced design (because of plant death after the start of the experiment) and low replication within families (N *=* 3 per family at beginning of experiment), the denominator degrees of freedom for fixed effects in the mixed models were calculated using the Kenward-Roger method (Kenward and Roger 1997).

## Results

### Plant damage

During the experiment, pollinators and herbivores visiting the *D. stramonium* plants included bees, *Manduca sexta* moths, syrphid flies, weevils, leaf miners, stink bugs, three-lined potato beetles (*Lema daturaphila*), and grasshoppers. Insects responsible for the most severe damage included flea beetles and *Manduca sexta. Manduca sexta* egg abundance during weekly leaf inspections peaked in late June and mid-August (17 eggs in the plot on 26 June 2014 and 8 eggs in the plot on 19 August 2014).

The level of herbivore damage increased over the field season, with JA induction providing initial protection that decreased over time and insecticide treatment protecting plants throughout the season. The repeated-measures analyses showed that the proportion of leaves damaged changed significantly over time *F*_1,636_ *=* 363.42, *P* < 0.01) with more damage occurring later in the season (Fig. 1). There was a significant main effect of JA induction treatment (*F*_1,550_ *=* 15.01, *P* < 0.01) but not of insecticide treatment (*F*_1,550_ *=* 2.07, *P =* 0.15) on plant damage, indicating the effect of JA induction early in the season (as detailed below). The effect of JA induction treatment and insecticide treatment (“INS”) changed over time (time*JA: *F*_1,636_ *=* 9.70, *P* < 0.01; time*INS: *F*_1,636_ *=* 11.47, *P* < 0.01). There were no strong differences in leaf damage among populations (*F*_2,316_ *=* 2.47, *P =* 0.09).

**Figure 1.**
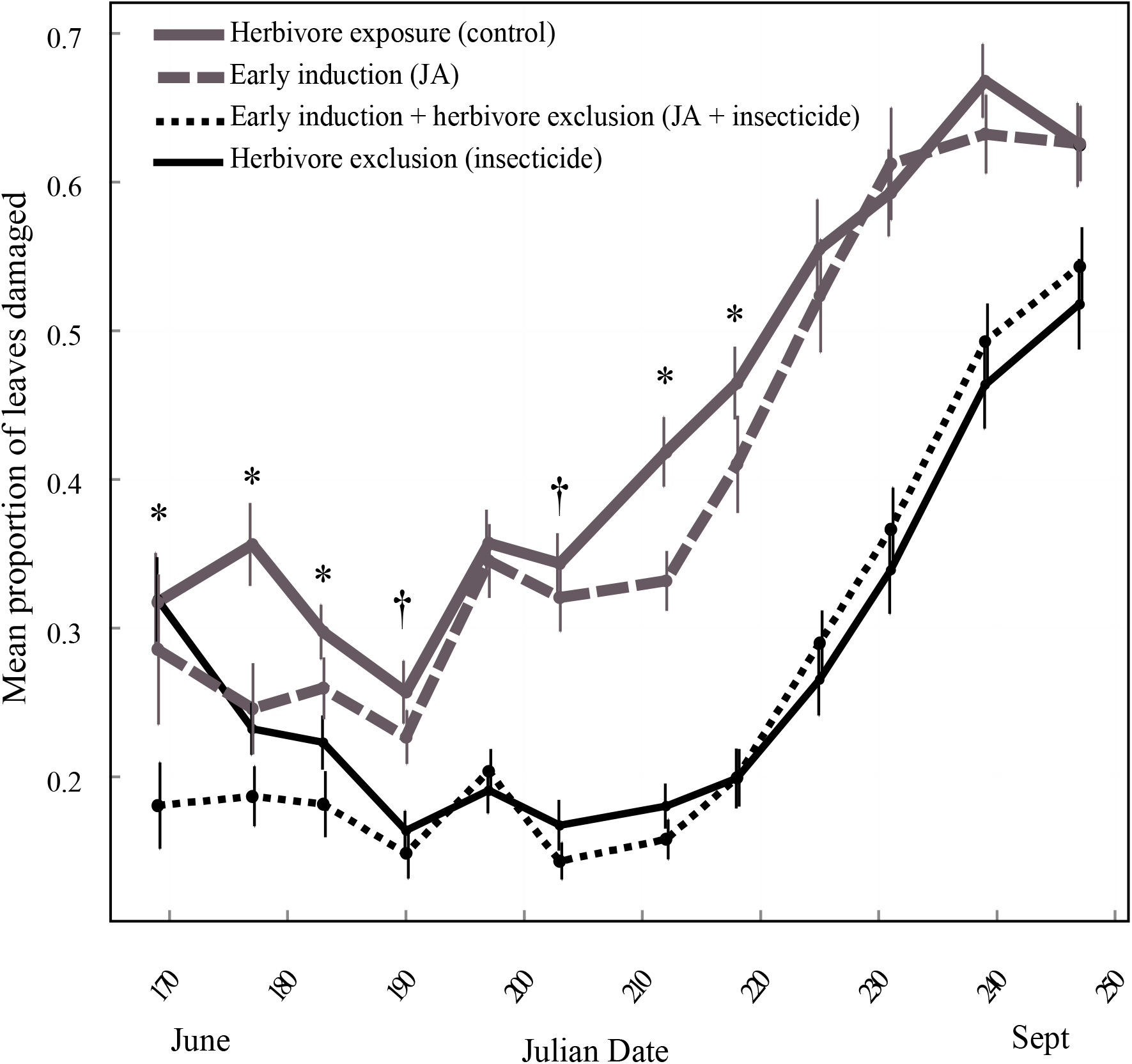
Mean proportion of leaves damaged per treatment over time. Vertical bars represent standard error. Insecticide treatment significantly decreased damage (P < 0.05) at each time point following the first date. Asterisks designate statistically significant reductions in damage for the jasmonic acid induction treatment based on mixed-effects models (statistical results in Table S1). * P < 0.05, † 0.05 < P < 0.1

Mixed-effects models for individual dates revealed that plants in the JA treatment (but not insecticide treatment) had a reduction in the proportion of leaves damaged beginning after only one week in the field (18 June 2014) (JA: *F*_1,142_ *=* 17.1, *P =* <0.01, INS: *F*_1,24.7_ *=* 0.49, *P =* 0.49) consistent with the main effect in the repeated measure analyses. Herbivore damage was significantly (or marginally significantly) lower for JA induction plants for the first 8 weeks in the field (except for week 5; 16 July 2014). There was no further effect of JA induction treatment during the last 4 weeks (13 August 2014 to 4 September 2014; when field estimates of damage were stopped) (Table S1). Weekly insecticide application significantly reduced the proportion of leaves damaged at each time point starting after two weeks in the field (26 June 2014) (*F*_1,159_ *=* 10.57, *P <* 0.01) and this reduction in damage was sustained throughout the season (Fig. 1, Table S1). Populations were marginally significantly different in the initial proportion of leaves damaged during the first week in the field (*F*_2,25.8_ *=* 2.99, *P =* 0.07) but populations were not consistently different in damage levels across the timepoints (Table S1).

### Plant size

The effect of the JA induction and insecticide treatments on plant size varied over the course of the season. Repeated-measures analyses showed that leaf numbers changed significantly over time (*F*_1,1960_ *=* 828.77, *P* < 0.01) and that the effect of JA induction treatment and insecticide treatment changed over time (time*JA: *F*_1,1960_ *=* 6.97, *P* < 0.01; time*INS: *F*_1,1960_ *=* 9.08, *P <* 0.01). There were also population-level size differences in leaf number (*F*_2,154_ *=* 3.65, *P =* 0.03), with plants in population W tending to have the most leaves while plants in population A tended to have the fewest leaves. Median leaf number at the final measurement (02 October 2014) was 210 leaves for population A (N = 57), 283 leaves for population H (N = 54), and 408 leaves for population W (N = 55).

Mixed-effects models for individual dates revealed reduced growth for plants in the JA induction treatment and plants exposed to herbivores (no-insecticide); however, the growth reduction associated with JA induction started earlier in the season than the growth reduction associated with herbivore exposure. The decrease in growth as a result of JA induction began during the second week in the field (26 June 2014); plants in this induced treatment had fewer leaves (*F*_1,151_ *=* 9.03, *P <* 0.01) and this significantly smaller size was sustained throughout the season. There was no difference in leaf number per plant between insecticide-treated plants and herbivore-exposed plants until the seventh week in the field (31 July 2014) when plants in the herbivore exposure treatment were marginally smaller (fewer leaves) than those receiving protection from herbivores (INS: *F*_1,19.7_ *=* 4.09, *P =* 0.06). Plant size in the herbivore exposed plants then remained significantly smaller throughout the experiment (Fig. 2, Table S1).

**Figure 2.**
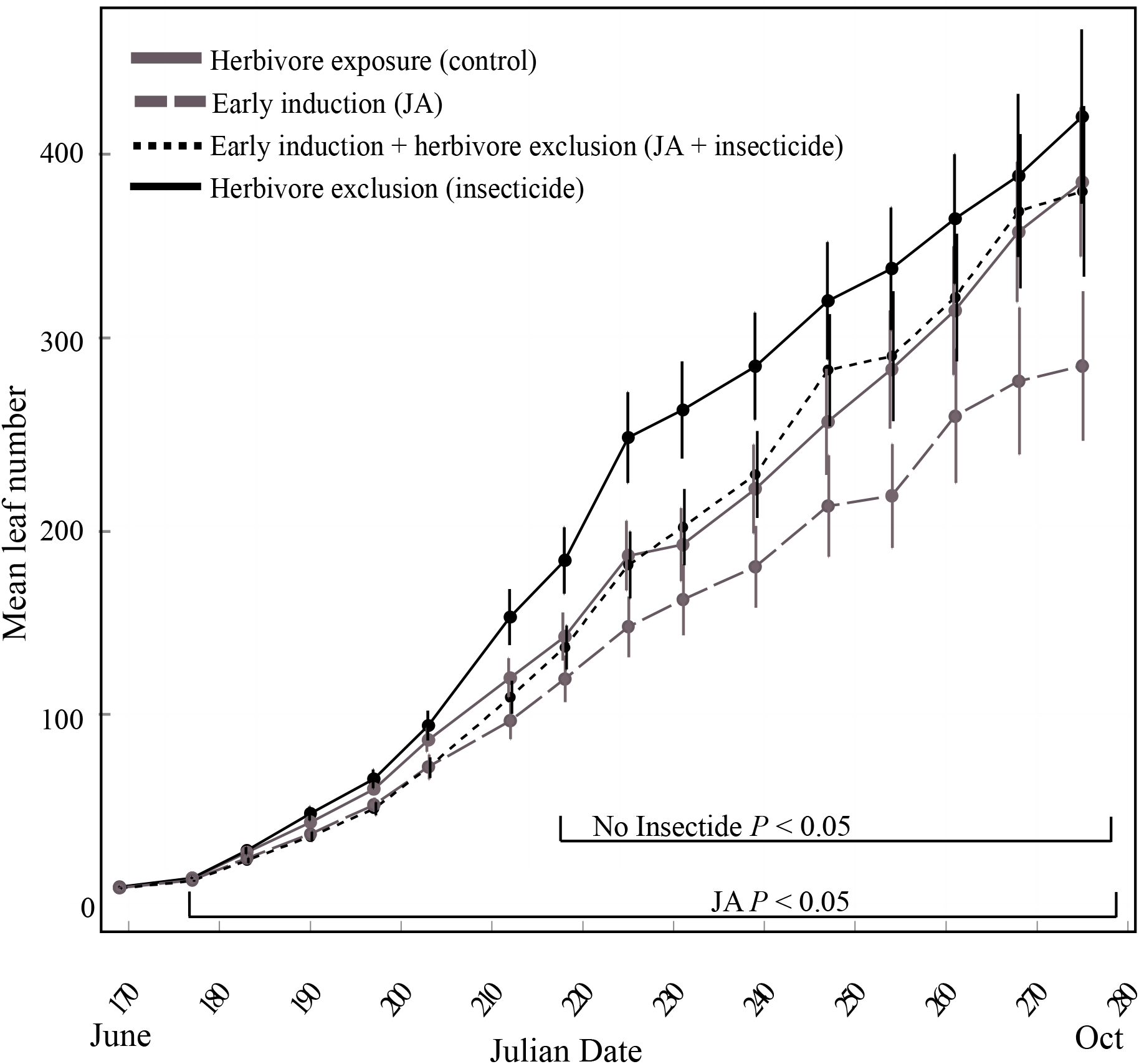
Mean leaf number per treatment over time. Vertical bars represent standard error. Brackets designate statistically significant reductions in leaf number for the jasmonic acid induction treatment or herbivore exposure (no insecticide) treatment based on mixed-effects models (statistical results in Table S1).

### Flowering time

Plants protected from herbivory with insecticide flowered later than plants fully exposed to herbivory (*F*_1,126_ *=* 25.51, *P* < 0.01), but flowering date was not affected by JA induction treatment (*F*_1,71.4_ *=* 0.73, *P =* 0.40) (Fig. 3). Flowering date did not differ among populations (*F*_2,11.5_ *=* 0.84, *P =* 0.46). Overall, median date of first flower was 14 July 2014 (N *=* 173, IQR *=* 10 July 2014-15 July 2014). Median date of the 23^rd^ flower produced (used for “late season” floral measurements) was 8 August 2014 (N *=* 139, IQR *=* 2 August 2014-24 August 2014).

**Figure 3.**
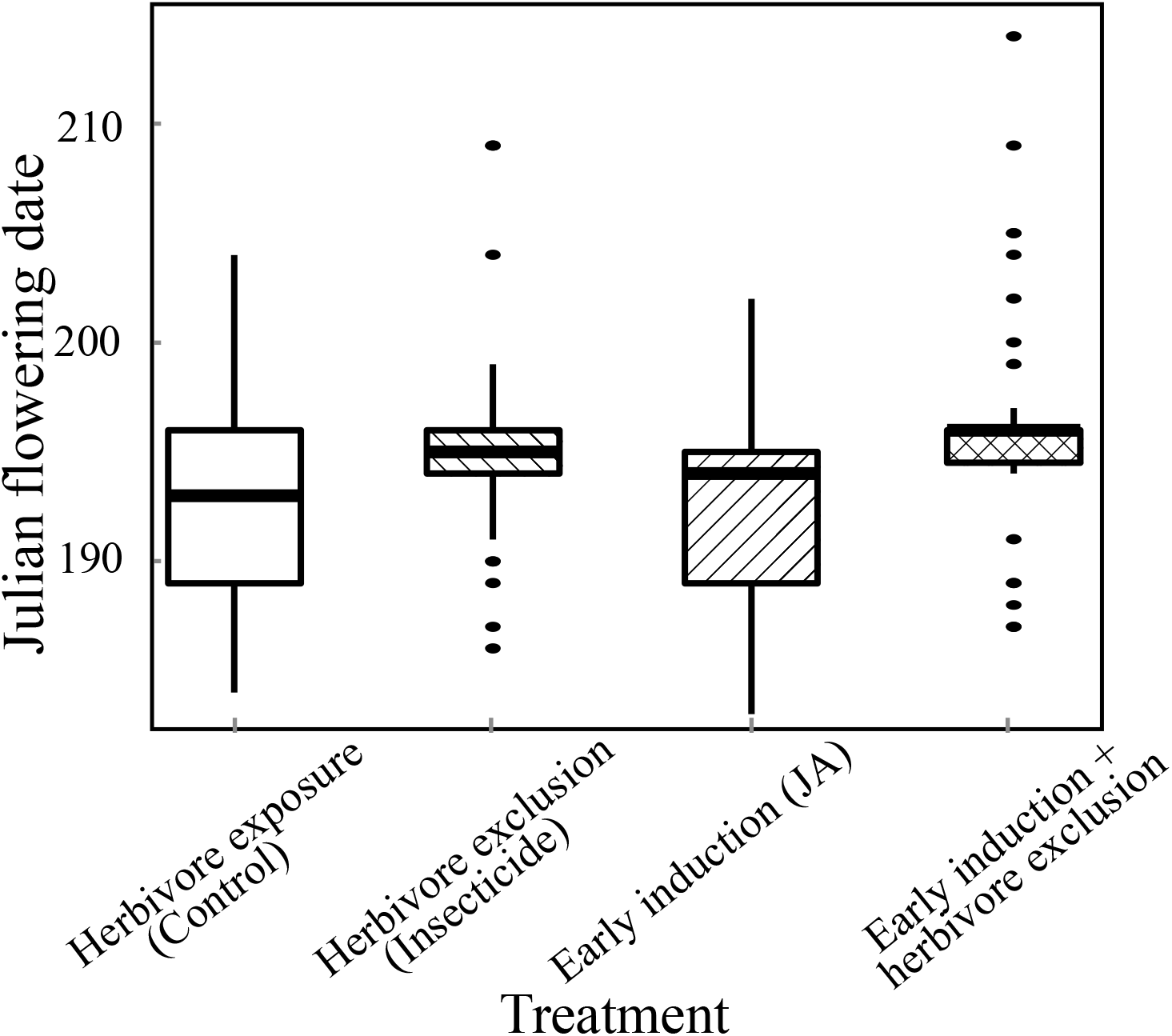
Date of first flowering for plants in each treatment. Plants exposed to natural herbivores (no insecticide) flowered earlier than plants treated with insecticide.

### Flower, fruit, and seed allocation

Early induction with JA and exposure to herbivores (no insecticide) reduced overall floral allocation. These plants produced fewer flowers (JA: *F*_1,129_ *=* 6.47, *P =* 0.01; INS: *F*_1,19.8_ *=* 5.71, *P =* 0.03). This reduction in flower number resulted in reduced fruit production for plants receiving the JA induction treatment (*F*_1,126_ *=* 6.74, *P =* 0.01) but not herbivore exposure plants (*F*_1,19.9_ *=* 3.61, *P =* 0.07). Total fruit production was an accurate assessment of total seed set because seed number per fruit did not differ among treatments or populations (JA: *F*_1,99.3_ = 1.35, *P* = 0.24; INS: *F*_1,85.9_ = 0.00, *P* = 0.95; POP: *F*_2,11.3_ = 2.5, *P* = 0.13). In addition to decreases in flower and fruit numbers, early floral size for JA induced plants was smaller than for those non-induced plants (*F*_1,120_ *=* 4.88, *P =* 0.03), but did not differ based on insecticide treatment (*F*_1,116_ *=* 1.37, *P =* 0.244). Later-season flowers did not differ significantly in size among treatments (JA: *F*_1,10_ *=* 1.95, *P =* 0.19; INS: *F*_1,110_ *=* 1.73, *P =* 0.19) but were smaller than early-season flowers (Table S2).

In addition to differences in floral allocation among treatments, there were population level differences in floral allocation. Population W produced the largest number of flowers (*F*_2,20.5_ *=* 5.09, *P =* 0.02) but Population A produced the largest flowers early in the season (*F*_2,123_ *=* 22.71, *P* < 0.01). Variation in fruit number among populations (*F*_2,20.5_ *=* 3.97, *P =* 0.03) mirrored the differences seen in flower numbers but not all flowers turned into successful fruits. Later-season flowers did not differ significantly in size among populations (*F*_2,12.2_ *=* 0.50, *P =* 0.62) (Table S2).

### Herkogamy

Late season flowers and flowers exposed to herbivores had herkogamy levels more conducive to selfing than outcrossing compared to early season flowers and flowers on plants in the insecticide treatment. In early flowers, anther-stigma overlap (a “selfing” morphology) was greater in herbivore exposed plants compared to insecticide-treated plants (*F*_1,18.7_ *=* 5.74, *P =* 0.03), but did not differ for JA induced and non-JA induced plants (*F*_1,18_ *=* 1.96, *P =* 0.18) or by population (*F*_2,21.7_ *=* 1.09, *P =* 0.35) (Fig. 4). Late-season flowers had more anther-stigma overlap than early-season flowers, with median herkogamy values below zero for all treatments (range -0.12 mm to -1.22 mm). However, herkogamy did not differ among treatments or populations late in the season (JA: *F*_1,66.1_ *=* 0.11, *P =* 0.75; INS: *F*_1,95.2_ *=* 0.51, *P =* 0.48; POP: *F*_2,11.8_ *=* 0.07, *P =* 0.93; JA*INS: *F*_1,95.2_ *=* 7.50, *P* < 0.01).

**Figure 4.**
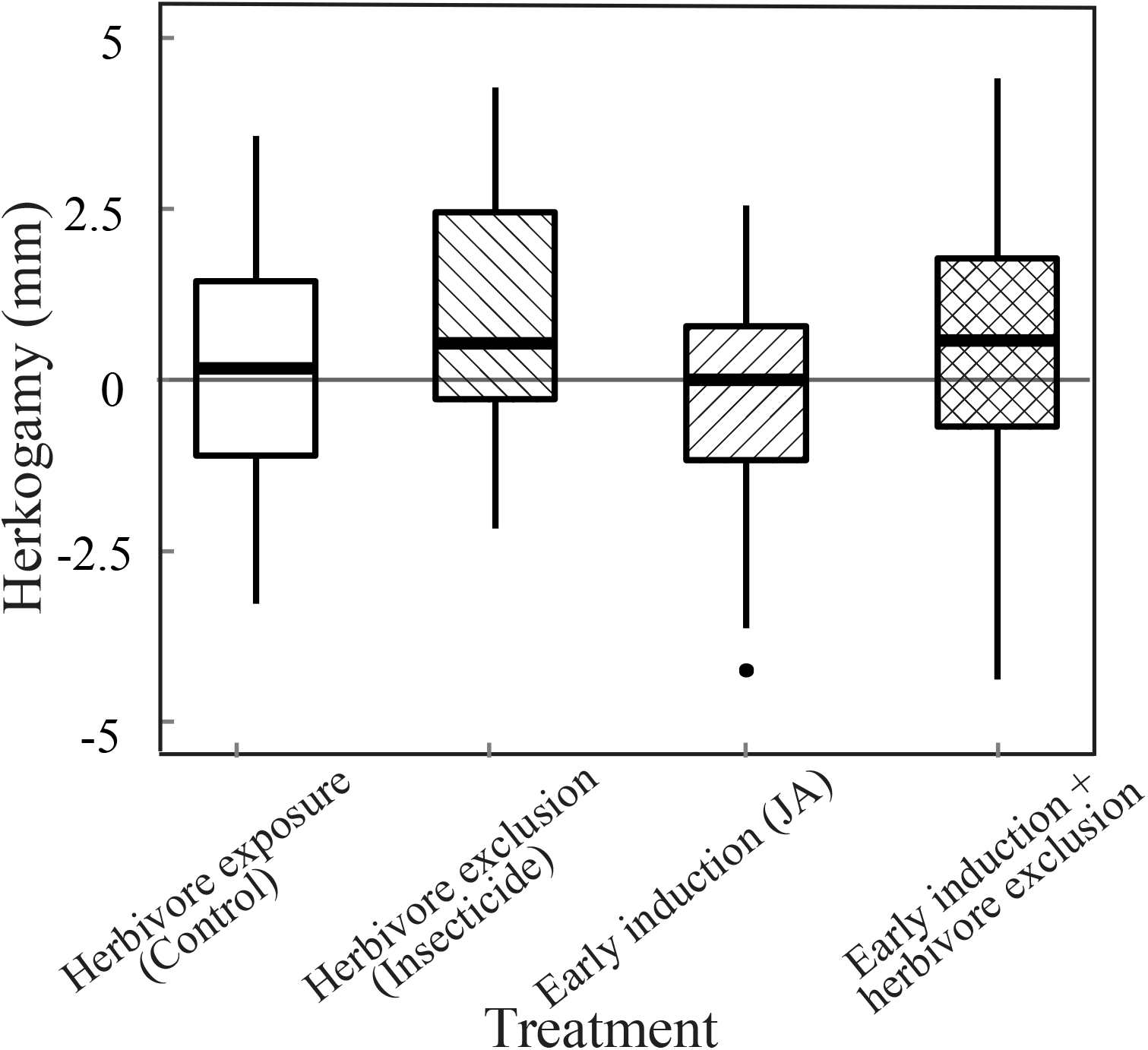
Herkogamy for flowers produced early in the season (1^st^-3^rd^ flowers per plant). Herkogamy (anther-stigma overlap) was greater in plants that were exposed to herbivores compared to plants in treatments that received insecticide.

## Discussion

Using a field experiment, I have shown that induced defenses are costly and result in changes in allocation to plant defense, plant growth, and reproduction over the lifetime of this annual plant. The cost of induced defense is revealed in plants that were induced early on with JA because even when these plants received low levels of damage in the field, they showed decreased plant size and floral allocation. The decrease in total fruit production associated with early JA induction indicates that the effects of early induction had long-lasting costs for fitness. Although plants without insecticide protection also had decreased flower numbers, fruit numbers did not differ. Abscission of developing fruits is common in *D. stramonium* and this difference in flower to fruit transitions among treatments likely indicates that smaller JA-induced plants did not have the resources to mature as many fruits.

One main caveat of field studies examining reduction of herbivory (via exclusion or insecticide) and defense induction is that it is very difficult to determine whether growth costs are the result of induction at a specific life stage or the result of the cumulative effect of long-term induction because even protected plants in the field receive some level of herbivore damage (Agrawal *et al*. 1999, Thaler 1999). However, examining the costs to reproductive allocation in the absence of herbivores is not likely to be biologically relevant, particularly when considering how plant defense, herbivory, and pollination interact to shape plant traits. In this study, plants were treated with JA prior to receiving field herbivory. Although all plants received some herbivory once in the field, plants that received JA had lower levels of damage early in the season but still showed lasting costs of induction.

The long-lasting effect of the jasmonic acid treatment on growth and reproduction may be the result of either the timing of induction or indirect effects of JA on defense. A plant’s ability to overcome damage may depend not only on the amount of signal received from herbivory (amount of damage) but also when this damage occurs. Young plants are thought to be more inducible than older plants (Karban and Baldwin 1997) but also have limited energetic resources because of a small photosynthetic area. This means that an early, transient signal of damage may be sufficient to induce defenses and the small size of the plant at induction may result in long-term growth costs. It is also possible that JA treatment changed initial herbivory in a way that altered the herbivore community throughout the season (Cippolini *et al*. 2003, Poelman and Kessler 2016, Visakorpi *et al*. 2019). Although costly, early defensive induction has been seen to increase early survival, which is crucial to plant establishment and reproduction (Baldwin 1988); this benefit of early induction may outweigh fitness losses when seedling/early juvenile herbivory is high.

This study is consistent with others showing decreased flower size and/or flower number following herbivore damage and/or induction. However, I found additional changes in floral morphology (herkogamy) that may alter interactions with pollinators apart from just overall reductions in floral allocation. Reduced vegetative and floral allocation are generally associated with reduced attractiveness to pollinators (reviewed in Johnson *et al*. 2015). This reduction in floral display would come at a cost to seed set unless plants have a way to compensate, such as by promoting self-fertilization when pollinators are absent/infrequent (Kessler and Halitschke 2009). In this study, plants exposed to herbivores (no insecticide) exhibited reductions in herkogamy, a trait that is correlated with selfing versus outcrossing rate (Motten and Stone 2000, Ivey and Carr 2005). This relationship between damage and herkogamy suggests that herbivory plays an important role in plant mating system dynamics (Campbell 2014). Decreased resources available to each sequential flower may partially explain the reduction in floral size and herkogamy in late season flowers, however a shift to this more “selfing” morphology may be adaptive as pollinators become less abundant later in the season (Brunet and Charlesworth 1995, Diggle 1995, Kliber and Eckert 2004).

Although a paternity analysis was not done for this experiment to confirm that herbivore damaged plants produced more selfed seed, prior studies in *D. stramonium* have shown lower herkogamy levels are associated with increased selfing for *D. stramonium* (Motten and Antonovics 1992, Stone and Motten 2002). The relationship between herkogamy and herbivore environment has also been explored in *Datura wrightii*, however the impact of herbivory on herkogamy was weak and dependent on interactions with water availability (Elle and Hare 2002). The stronger response in *D. stramonium* is likely explained by the tighter association between herkogamy and outcrossing rates in *D. stramonium* compared to *D. wrightii* (Elle and Hare 2002). This is consistent with the prediction that the ecological interactions between herbivory, pollination, and inbreeding depression may maintain mixed mating as a stable strategy, because the rates and benefits of selfing and outcrossing vary depending on environmental influences (Goodwillie *et al*. 2005, Campbell *et al*. 2013, Carr and Eubanks 2014).

Plants fully exposed to herbivores (no insecticide) flowered earlier in the season indicating that herbivore environment can change flowering phenology, thereby altering the timing of floral availability to pollinators and potential mates. In other systems, the effect of herbivory on flowering time varies, with some systems showing early flowering in response to herbivory (Hoffmeister *et al*. 2015) and some showing delayed flowering (Strauss *et al*. 1996; Hanley and Fegan 2007). While the direction of the shift in flowering date likely depends on a variety of factors including plant resource status and life history, herbivore-induced shifts in flowering time change the pollinator and mate environment that undamaged and damaged plants experience. A change in the timing of flowering can also impact fitness as a result of abiotic conditions. Plants with later flowering dates may have fruits that are still maturing later in the season and at risk of frost damage/loss. Many of the plants in this study were still developing fruits at the timing of the first hard frost in November, which resulted in the loss of those seeds.

Overall this study demonstrates that early defensive induction and herbivory result in long-lasting effects on plant growth and reproduction. The magnitude of the effect of these leaf and floral traits on plant fitness depends on abiotic and biotic environmental conditions. These changes may have implications not only for individual plant fitness, but also for interactions among plants (by altering mating environment) and for interactions with pollinators (via changes in total floral allocation and floral phenology).

## Acknowledgements

N. Harby and the Ralph M. Kriebe Herbarium at Purdue University supplied the seeds from the Wabash and US Hwy 53 populations. D. Haak supplied the seeds from the Applegate population and provided guidance on experimental methods. L. Delph provided feedback on the experimental design and writing. J. Bever, C. Lively, L. Moyle, and R. Raguso gave constructive comments on the manuscript and J. Bever and D. Castillo provided statistical help. A. Bouwkamp, J. Greenburg, and C. King assisted with planting and data collection and the Indiana University Greenhouses aided in field setup. This material is based upon work supported by the National Science Foundation Graduate Research Fellowship.

## Data accessibility

Data to be deposited in the Dryad repository.

## Supplementary information

**Figure S1**. Layout of field plot in Bloomington, Indiana showing plants (denoted with “X”), treatments, and plots within common garden. Green highlighted cells indicate jasmonic acid early induction treatment while yellow highlighted cells indicate control (EtOH in water) treatment. Orange outlined cells indicate weekly insecticide spray while blue outlined cells indicate weekly control (water) spray (herbivore exposure).

**Table S1**. Statistical table for Box-Cox transformations and mixed-effects models for leaf number and proportion of leaves damaged at each time point. Bold font indicates significance at *P* < 0.05 and underline indicates marginal significance at *P* < 0.10.

**Table S2**. Population-level plant reproductive measurements among jasmonic-acid and herbivore-induced defense treatments. Corolla length is averaged over the 1^st^-3^rd^ or 23^rd^-25^th^ flowers open per plant for the early and late measures, respectively. Median values (followed by sample size and interquartile range) are given for each population separately and combined (“overall”). Treatments are: control/none (“C”), insecticide (“I”), jasmonic acid early induction (“J”) and jasmonic acid early induction plus herbivore exposure via insecticide (“J+I”).

## Notes

### Competing Interest Statement

The authors have declared no competing interest.

