## Supplementary material for "Costly induction of defense reduces plant growth and alters reproductive traits in mixed-mating *Datura stramonium*": Figure S1

Deidra J. Jacobsen

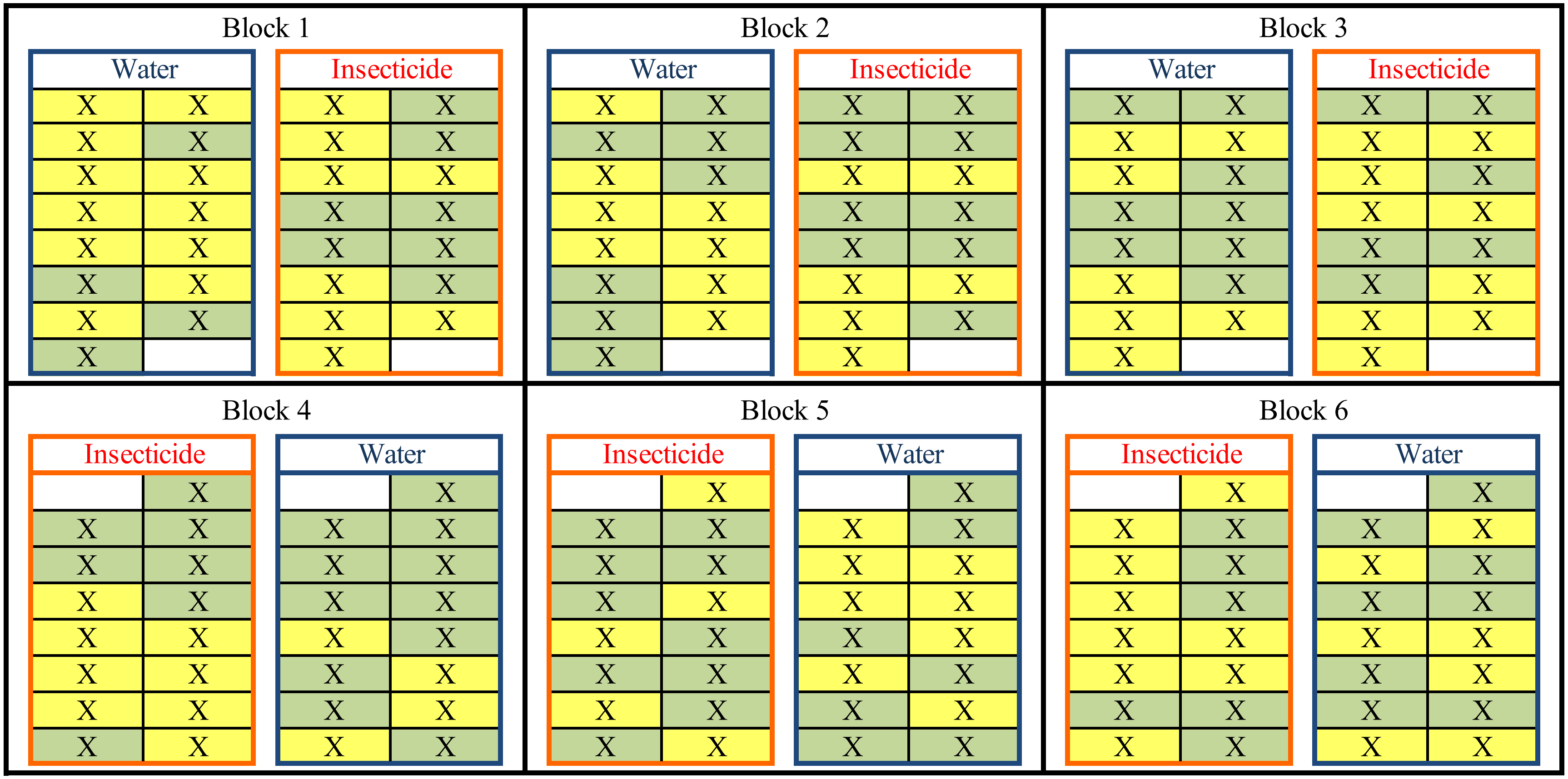


Figure S1. Layout of field plot in Bloomington, Indiana showing plants (denoted with “X”), treatments, and blocks within common garden. Green highlighted cells indicate jasmonic acid early induction treatment while yellow highlighted cells indicate control (EtOH in water) treatment. Orange outlined cells indicate weekly insecticide spray (herbivore exclusion) while blue outlined cells indicate weekly control (water) spray (herbivore exposure).
