## Supplementary material for "Costly induction of defense reduces plant growth and alters reproductive traits in mixed-mating *Datura stramonium*": Table S1

| Leaf numbers | | | JA | | | INS | | | JA*INS | | | POP | | | POP*INS | | | POP*JA | | | POP*INS*JA | | |
|---|---|---|---|---|---|---|---|---|---|---|---|---|---|---|---|---|---|---|---|---|---|---|---|
| Date | Box-Cox | N | df | F | P | df | F | P | df | F | P | df | F | P | df | F | P | df | F | P | df | F | P |
| 29-May | 1.4 | 178 | 1, 154 | 1.64 | 0.20 | 1, 136 | 0.33 | 0.57 | 1, 154 | 0.64 | 0.43 | 2, 11.9 | 7.74 | **0.01** | 2, 135 | 0.19 | 0.83 | 2, 154 | 1.09 | 0.34 | 2, 150 | 0.59 | 0.56 |
| 10-Jun | 1.4 | 178 | 1, 11.5 | 2.26 | 0.16 | 1, 136 | 0.34 | 0.56 | 1, 136 | 0.09 | 0.77 | 2, 11.4 | 13.94 | **0.00** | 2, 136 | 0.07 | 0.94 | 2, 11.5 | 1.19 | 0.34 | 2, 136 | 0.47 | 0.63 |
| 18-Jun | 1.6 | 178 | 1, 12.1 | 2.19 | 0.16 | 1, 24.7 | 0.01 | 0.92 | 1, 24.7 | 3.51 | 0.07 | 2, 11.9 | 6.07 | **0.02** | 2, 24.7 | 0.11 | 0.90 | 2, 12.1 | 0.75 | 0.50 | 2, 24.7 | 0.05 | 0.95 |
| 26-Jun | -0.2 | 176 | 1, 151 | 9.03 | **0.00** | 1, 152 | 0.47 | 0.49 | 1, 151 | 0.17 | 0.68 | 2, 10.9 | 5.66 | **0.02** | 2, 152 | 0.96 | 0.39 | 2, 151 | 1.43 | 0.24 | 2, 151 | 0.04 | 0.96 |
| 2-Jul | 0.3 | 174 | 1, 143 | 10.73 | **0.00** | 1, 63.7 | 0.16 | 0.69 | 1, 143 | 0.79 | 0.38 | 2, 11.5 | 4.36 | **0.04** | 2, 63.3 | 0.99 | 0.38 | 2, 141 | 1.00 | 0.37 | 2, 140 | 0.83 | 0.44 |
| 9-Jul | 0.4 | 174 | 1, 131 | 14.58 | **0.00** | 1, 79.7 | 0.19 | 0.67 | 1, 131 | 1.13 | 0.29 | 2, 11.7 | 3.46 | 0.07 | 2, 81.8 | 0.74 | 0.48 | 2, 128 | 0.60 | 0.55 | 2, 126 | 3.29 | **0.04** |
| 16-Jul | 0.6 | 174 | 1, 123 | 12.31 | **0.00** | 1, 10.4 | 0.13 | 0.73 | 1, 123 | 1.37 | 0.24 | 2, 11.6 | 4.19 | **0.04** | 2, 10.5 | 0.20 | 0.82 | 2, 120 | 0.59 | 0.56 | 2, 119 | 3.43 | 0.36 |
| 22-Jul | 0.5 | 173 | 1, 127 | 9.93 | **0.00** | 1, 10.4 | 0.47 | 0.51 | 1, 127 | 0.4 | 0.53 | 2, 11.3 | 4.85 | **0.03** | 2, 10.4 | 0.26 | 0.78 | 2, 124 | 0.91 | 0.41 | 2, 124 | 5.49 | **0.01** |
| 31-Jul | 0.2 | 171 | 1, 130 | 9.65 | **0.00** | 1, 19.7 | 4.09 | 0.06 | 1, 130 | 0.16 | 0.69 | 2, 19.9 | 6.53 | **0.01** | 2, 19.9 | 0.14 | 0.87 | 2, 127 | 1.13 | 0.33 | 2, 128 | 4.77 | **0.01** |
| 6-Aug | 0.3 | 171 | 1, 130 | 7.84 | **0.01** | 1, 20.1 | 4.44 | **0.05** | 1, 131 | 0.37 | 0.55 | 2, 20.5 | 6.33 | **0.01** | 2, 20.3 | 0.24 | 0.79 | 2, 128 | 1.20 | 0.31 | 2, 128 | 4.93 | **0.01** |
| 13-Aug | 0.3 | 171 | 1, 132 | 9.91 | **0.00** | 1, 19.7 | 7.57 | **0.01** | 1, 132 | 0.52 | 0.47 | 2, 20.2 | 7.90 | **0.00** | 2, 20.1 | 0.17 | 0.84 | 2, 130 | 0.78 | 0.46 | 2, 130 | 5.25 | **0.01** |
| 19-Aug | 0.3 | 171 | 1, 130 | 7.01 | **0.01** | 1, 19.9 | 9.62 | **0.01** | 1, 131 | 0.61 | 0.44 | 2, 20.5 | 9.29 | **0.00** | 2, 20.3 | 0.02 | 0.98 | 2, 128 | 0.78 | 0.46 | 2, 128 | 5.53 | **0.01** |
| 27-Aug | 0.3 | 171 | 1, 130 | 7.74 | **0.01** | 1, 19.7 | 9.58 | **0.01** | 1, 131 | 0.12 | 0.73 | 2, 20.4 | 8.24 | **0.00** | 2, 20.2 | 0.05 | 0.95 | 2, 128 | 1.01 | 0.37 | 2, 128 | 4.84 | 0.10 |
| 4-Sep | 0.4 | 171 | 1, 128 | 4.47 | **0.04** | 1, 20 | 13.37 | **0.00** | 1, 129 | 0.02 | 0.90 | 2, 20.7 | 6.64 | **0.01** | 2, 20.5 | 0.08 | 0.93 | 2, 126 | 0.35 | 0.70 | 2, 126 | 5.46 | **0.01** |
| 11-Sep | 0.3 | 170 | 1, 125 | 7.04 | **0.01** | 1, 20.9 | 10.46 | **0.00** | 1, 125 | 0.07 | 0.79 | 2, 21.5 | 4.36 | **0.03** | 2, 21.4 | 0.16 | 0.85 | 2, 123 | 0.41 | 0.67 | 2, 122 | 4.44 | **0.01** |
| 18-Sep | 0.4 | 168 | 1, 19.8 | 5.32 | **0.03** | 1, 94.8 | 8.50 | **0.00** | 1, 116 | 0.06 | 0.80 | 2, 25.8 | 4.09 | **0.03** | 2, 9,9 | 0.05 | 0.95 | 2, 20.1 | 0.42 | 0.66 | 2, 113 | 3.21 | **0.04** |
| 25-Sep | 0.4 | 167 | 1, 137 | 5.70 | **0.02** | 1, 93.6 | 7.91 | **0.01** | 1, 137 | 0.93 | 0.34 | 2, 99.9 | 3.11 | **0.05** | 2, 96.2 | 0.05 | 0.96 | 2, 135 | 0.41 | 0.67 | 2, 133 | 1.72 | 0.18 |
| 2-Oct | 0.4 | 166 | 1, 128 | 6.96 | **0.01** | 1, 21.3 | 5.87 | **0.00** | 1, 128 | 0.88 | 0.35 | 2, 22.1 | 3.35 | 0.05 | 2, 21.9 | 0.03 | 0.97 | 2, 127 | 0.22 | 0.80 | 2, 126 | 3.18 | **0.05** |

| Proportion leaves damaged | | | JA | | | INS | | | JA*INS | | | POP | | | POP*INS | | | POP*JA | | | POP*INS*JA | | |
|---|---|---|---|---|---|---|---|---|---|---|---|---|---|---|---|---|---|---|---|---|---|---|---|
| Date | Box-Cox | N | df | F | P | df | F | P | df | F | P | df | F | P | df | F | P | df | F | P | df | F | P |
| 18-Jun | (+1) -0.7 | 178 | 1, 142 | 17.07 | **0.00** | 1, 24.7 | 0.49 | 0.49 | 1, 142 | 0.75 | 0.39 | 2, 25.8 | 2.99 | 0.07 | 2, 25.3 | 0.10 | 0.90 | 2, 142 | 0.27 | 0.77 | 2, 144 | 0.69 | 0.50 |
| 26-Jun | (+1) -1.5 | 176 | 1, 164 | 13.08 | **0.00** | 1, 159 | 10.57 | **0.00** | 1, 164 | 1.58 | 0.21 | 2, 160 | 0.17 | 0.84 | 2, 163 | 0.09 | 0.92 | 2, 163 | 1.74 | 0.18 | 2, 159 | 0.09 | 0.91 |
| 2-Jul | (+1) -1.6 | 174 | 1, 162 | 5.99 | **0.02** | 1, 162 | 17.14 | **0.00** | 1, 162 | 0.10 | 0.76 | 2, 162 | 0.17 | 0.85 | 2, 162 | 1.07 | 0.35 | 2, 162 | 1.54 | 0.22 | 2, 162 | 0.38 | 0.68 |
| 9-Jul | (+1) -2.7 | 174 | 1, 136 | 2.86 | 0.09 | 1, 20 | 23.72 | **0.00** | 1, 136 | 0.00 | 0.99 | 2, 20.1 | 0.00 | 1.00 | 2, 135 | 0.90 | 0.41 | 2, 135 | 0.90 | 0.41 | 2, 136 | 0.79 | 0.46 |
| 16-Jul | (+1) -2.1 | 174 | 1, 68.4 | 0.11 | 0.74 | 1, 126 | 72.21 | **0.00** | 1, 125 | 1.28 | 0.26 | 2, 12.5 | 0.68 | 0.52 | 2, 125 | 1.47 | 0.23 | 2, 69.4 | 0.45 | 0.64 | 2, 123 | 0.67 | 0.51 |
| 22-Jul | 0.4 | 173 | 1, 146 | 3.48 | 0.06 | 1, 144 | 126.3 | **0.00** | 1, 145 | 0.00 | 1.00 | 1, 12 | 0.44 | 0.65 | 2, 145 | 2.90 | 0.06 | 2, 146 | 2.58 | 0.08 | 2, 147 | 1.41 | 0.25 |
| 31-Jul | (+1) -1.4 | 171 | 1, 155 | 11.41 | **0.00** | 1, 154 | 170.5 | **0.00** | 1, 155 | 1.61 | 0.21 | 2, 156 | 1.77 | 0.17 | 2, 155 | 0.36 | 0.70 | 2, 155 | 0.59 | 0.55 | 2, 156 | 2.11 | 0.13 |
| 6-Aug | 0.4 | 171 | 1, 141 | 4.16 | **0.04** | 1, 59.8 | 155.3 | **0.00** | 1, 142 | 1.67 | 0.20 | 2, 12 | 7.60 | **0.01** | 2, 60.5 | 1.68 | 0.20 | 2, 140 | 1.32 | 0.27 | 2, 141 | 2.11 | 0.13 |
| 13-Aug | 0.4 | 171 | 1, 41.7 | 0.22 | 0.64 | 1, 47.4 | 88.20 | **0.00** | 1, 41.4 | 1.53 | 0.22 | 2, 50 | 3.35 | **0.04** | 2, 48.7 | 0.49 | 0.62 | 2, 41.4 | 1.75 | 0.19 | 2, 42.8 | 1.44 | 0.25 |
| 19-Aug | 0.6 | 171 | 1, 18.6 | 0.03 | 0.86 | 1, 29.4 | 63.92 | **0.00** | 1, 18.8 | 0.04 | 0.85 | 2, 30.2 | 1.75 | 0.19 | 2, 30.1 | 1.97 | 0.16 | 2, 18.5 | 1.46 | 0.26 | 2, 19.3 | 1.71 | 0.21 |
| 27-Aug | 0.9 | 171 | 1, 23.3 | 0.14 | 0.72 | 1, 134 | 48.70 | **0.00** | 1, 136 | 1.44 | 0.23 | 2, 23.9 | 0.46 | 0.64 | 2, 136 | 0.55 | 0.58 | 2, 23.3 | 1.34 | 0.28 | 2, 137 | 1.94 | 0.15 |
| 4-Sep | 1.1 | 171 | 1, 146 | 0.14 | 0.71 | 1, 145 | 14.07 | **0.00** | 1, 146 | 0.12 | 0.73 | 2, 12.5 | 1.15 | 0.35 | 2, 146 | 0.20 | 0.78 | 2, 146 | 0.13 | 0.88 | 2, 147 | 1.06 | 0.35 |

**Table S1.** Statistical table for Box-Cox transformations and mixed-effects models for leaf number and proportion of leaves damaged at each time point. Bold font indicates significance at $P < 0.05$ and underline indicates marginal significance at $P < 0.10$.
