## Supplementary material for "Costly induction of defense reduces plant growth and alters reproductive traits in mixed-mating *Datura stramonium*": Table S2

| | | Population | | | Overall |
|---|---|---|---|---|---|
| Trait | Treatment | Applegate (A) | Highway 52 (H) | Wabash (W) | |
| Early corolla length (cm) | C | 10.80 (15, 10.52-11.18) | 10.98 (15, 10.76-11.08) | 10.55 (15, 10.29-10.92) | 10.80 (45, 10.43-11.06) |
| | I | 11.23 (15, 11.06-11.41) | 10.72 (13, 10.6-11.09) | 10.68 (12, 10.48-10.92) | 10.89 (40, 10.62-11.24) |
| | J | 10.84 (15, 10.82-11.12) | 10.46 (15, 10.42-10.68) | 10.59 (15, 10.22-10.82) | 10.68 (45, 10.42-10.68) |
| | J+I | 11.10 (14, 10.76-11.56) | 10.70 (14, 10.43-11.05) | 10.32 (15, 10.24-10.72) | 10.74 (43, 10.32-11.08) |
| Late corolla length (cm) | C | 9.71 (13, 7.78-10.75) | 10.08 (11, 9.78-10.86) | 9.64 (14, 9.39-10.16) | 9.83 (38, 9.37-10.5) |
| | I | 9.91 (13, 9.60-11.01) | 10.11 (11, 9.57-10.82) | 10.38 (12, 9.61-10.52) | 10.23 (36, 9.60-10.75) |
| | J | 10.01 ( 9, 9.34-10.39) | 9.79 ( 7, 9.43-10.38) | 9.65 (15, 9.08-10.00) | 9.73 (31, 9.30-10.13) |
| | J+I | 10.03 (10, 9.29-10.15) | 9.99 (13, 9.19-10.49) | 9.62 (13, 9.07- 9.64) | 9.95 (36, 9.19-10.39) |
| Total flower number per plant | C | 30 (15, 26-73) | 51 (15, 30.5-71.5) | 73 (15, 44.5-83.5) | 51 (45, 29.00-80.00) |
| | I | 62 (15, 36.5-82) | 42 (13, 27-70) | 90 (12, 61.3-98.3) | 62 (40, 40.25-89.5) |
| | J | 33.5 (14, 19-44.8) | 29 (14, 15.25-49.5) | 60 (15, 49.5-96) | 43 (43, 19.50-59.00) |
| | J+I | 45.5 (14, 23-56) | 52 (14, 32-74.25) | 52.5 (14, 41.5-64) | 49.5 (42, 31.50-71.25) |
| Total fruit number per plant | C | 20 (15, 15.5-47.5) | 33 (15, 18-57.5) | 52 (15, 24.5-58.5) | 33 (45, 16.00-57.00) |
| | I | 35 (15, 19.5-56) | 29 (13, 19-41) | 49.5 (12, 34.5-59.8) | 37 (40, 22.00-55.50) |
| | J | 24.5 (14, 13.3-30) | 18.5 (14, 11.3-30.8) | 44 (15, 29-59.5) | 27 (43, 13.50-41.50) |
| | J+I | 29.5 (14, 13-34) | 36 (14, 20.3-48) | 35 (14, 25.5-40) | 32 (42, 18.50-41.50) |

**Table S2**. Population-level plant reproductive measurements among jasmonic-acid and herbivore-induced defense treatments. Corolla length is averaged over the 1$^{st}$-3$^{rd}$ or 23$^{rd}$-25$^{th}$ flowers open per plant for the early and late measures, respectively. Median values (followed by sample size and interquartile range) are given for each population separately and combined ("overall"). Treatments are: control/none ("C"), insecticide (herbivore exclusion) ("I"), jasmonic acid early induction ("J") and jasmonic acid early induction plus herbivore exposure via insecticide ("J+I").
